# cGAS is predominantly a nuclear protein

**DOI:** 10.1101/486118

**Authors:** Hannah E. Volkman, Stephanie Cambier, Elizabeth E. Gray, Daniel B. Stetson

**Affiliations:** Department of Immunology, University of Washington School of Medicine, 750 Republican St, Seattle, WA 98109 USA

## Abstract

cGAS is an intracellular innate immune sensor that detects double-stranded DNA. The presence of billions of base pairs of genomic DNA in all nucleated cells raises the question of how cGAS is not constitutively activated. A widely accepted explanation for this is the sequestration of cGAS in the cytosol, which is thought to prevent cGAS from accessing nuclear DNA. Here, we demonstrate that cGAS is predominantly a nuclear protein, regardless of cell cycle phase or cGAS activation status. We show that nuclear cGAS is tethered tightly by a salt-resistant interaction. This tight tethering is independent of the domains required for cGAS activation, and it requires intact nuclear chromatin. We propose that tethering prevents activation of cGAS by genomic DNA, and that it might enable cGAS to distinguish between self DNA and foreign DNA within the nucleus.

## Introduction

The cGAS-STING DNA sensing pathway has emerged as a key innate immune response that is important for antiviral immunity (Goubau et al., 2013), contributes to specific autoimmune diseases (Crowl et al., 2017), and mediates important aspects of antitumor immunity (Li and Chen, 2018). cGAS binds to double-stranded DNA and catalyzes the formation of cyclic GMP-AMP (Sun et al., 2013; Wu et al., 2013), a diffusible cyclic dinucleotide that activates the endoplasmic adaptor protein STING (Ishikawa and Barber, 2008). Activated STING then serves as a platform for the inducible recruitment of the TBK1 kinase, which phosphorylates and activates the transcription factor IRF3, leading to the induction of the type I interferon mediated antiviral response (Liu et al., 2015).

cGAS is important for “cytosolic DNA sensing,” a term that was first proposed in 2006, years before the discovery of STING and cGAS as the essential adaptor and unique sensor of this pathway (Stetson and Medzhitov, 2006). At the time, a key conundrum was how a sequence-independent DNA sensing pathway that was activated by the sugar-phosphate backbone of DNA could avoid constant autoreactivity against genomic DNA that is present in all nucleated cells. We proposed the possibility that the sensor would be sequestered in the cytosol, separated by the nuclear envelope from genomic DNA, and that the inappropriate appearance of DNA in the cytosol would enable detection of foreign DNA while maintaining “ignorance” to self DNA (Stetson and Medzhitov, 2006). The discovery of cGAS and cGAMP, and subsequent structural studies of cGAS binding to DNA provided an elegant explanation for the sequence-independence of the response, the requirement for double-stranded DNA as a ligand, the contribution of the deoxyribose sugar-phosphate backbone of DNA to detection, and the definitive link between DNA sensing and STING (Civril et al., 2013; Li et al., 2013; Sun et al., 2013; Wu et al., 2013). However, the precise location of cGAS prior to its activation has remained largely unexplored, in part because of a lack of tools to track endogenous cGAS. Cytosolic DNA sensing has persisted as the mechanistic framework that guides the field.

There are a number of important problems with the model of cytosolic DNA sensing. First, nearly all DNA viruses (with the exception of the poxviruses) replicate their DNA exclusively in the nucleus. Studies of cGAS-deficient mouse and human cells have revealed that cGAS is important for the IFN-mediated antiviral response to these nuclear-replicating viruses, including herpesviruses (Ma et al., 2015; Wu et al., 2015). Moreover, retroviruses and lentiviruses are detected by cGAS (Gao et al., 2013; Lahaye et al., 2013; Rasaiyaah et al., 2013), but they shield their DNA within a capsid during reverse transcription in the cytosol, releasing this DNA for integration into the genome upon translocation into the nucleus. To fit the concept of cytosolic DNA sensing, current models envision that such viruses “leak” DNA into the cytosol during cellular entry or during exit. Second, as noted in the original description of cytosolic DNA sensing (Stetson and Medzhitov, 2006), cell division results in the breakdown of the nuclear envelope and the mixing of cytosolic and nuclear contents, which challenges a simple model of cytosolic sequestration as the basis for self/non-self discrimination by cGAS. Indeed, recent studies have demonstrated that cGAS can be found associated with mitotic chromosomes (Yang et al., 2017). This association is thought to be mediated by the generic DNA binding properties of cGAS, and it has been proposed that upon resolution of cell division and reformation of the nuclear envelope, cGAS is redistributed to the cytosol (Yang et al., 2017).

Here, we use confocal microscopy and biochemical characterization to determine the resting localization of endogenous cGAS prior to activation. We unexpectedly find that the vast majority of cGAS is in the nucleus, regardless of whether cells are rapidly dividing or post-mitotic. Moreover, we demonstrate that cGAS is tethered tightly in the nucleus by a salt-resistant interaction that rivals that of histones in its strength. We propose that this tight tethering explains the fact that nuclear cGAS is not constitutively activated by self DNA.

## Results

### Endogenous cGAS is predominantly a nuclear protein

We screened numerous commercially available antibodies to human cGAS for their ability to identify endogenous cGAS unambiguously and specifically using immunofluorescence microscopy. We chose to image HeLa cells, which express endogenous cGAS that is inactive in resting cells and potently activated to produce cGAMP upon transfection of calf thymus DNA (Figure 1 – Figure supplement 1A). Despite this potent activation of cGAS and production of cGAMP after DNA transfection, HeLa cells fail to activate the type I interferon response because the E7 oncoprotein of human papillomavirus 18 blocks STING-dependent signaling (Lau et al., 2015). To test for specificity of staining, we used lentiCRISPR to generate clonal lines of cGAS-deficient HeLa cells (Figure 1 – Figure supplement 1B; (Gray et al., 2016). We found that the D1D3G rabbit monoclonal antibody that detects an epitope in the N-terminus of human cGAS was suitable for analysis of endogenous cGAS by microscopy. Unexpectedly, endogenous cGAS was localized almost exclusively in the nuclei of all HeLa cells, with little cytosolic staining (Figure 1A). Identically prepared cGAS-deficient HeLa cells had no detectable immunostaining, confirming the specificity of this antibody for endogenous cGAS (Figure 1A). We noted three additional reproducible patterns of cGAS localization in addition to the uniform nuclear staining. First, as obesrved previously (Yang et al., 2017), we found that cGAS was associated with condensed mitotic chromatin (Figure 1B). Second, we found cGAS in rare, spontaneous, DAPI-positive, micronucleus-like extranuclear structures (Figure 1B). Whereas cGAS localization to micronuclei has been reported recently in a number of studies that primarily visualized overexpressed cGAS (Bartsch et al., 2017; Dou et al., 2017; Gluck et al., 2017; Harding et al., 2017; Mackenzie et al., 2017; Yang et al., 2017), we found that all cells with such structures also had extensive endogenous cGAS staining in the main nucleus (Figure 1B). Third, we found endogenous cGAS localized to “chromatin bridges” between adjacent cells (Figure 1B), the origins of which are thought to involve chromosome fusions and incomplete segregation of DNA between daughter cells during mitosis (Maciejowski et al., 2015).

**Figure 1:**
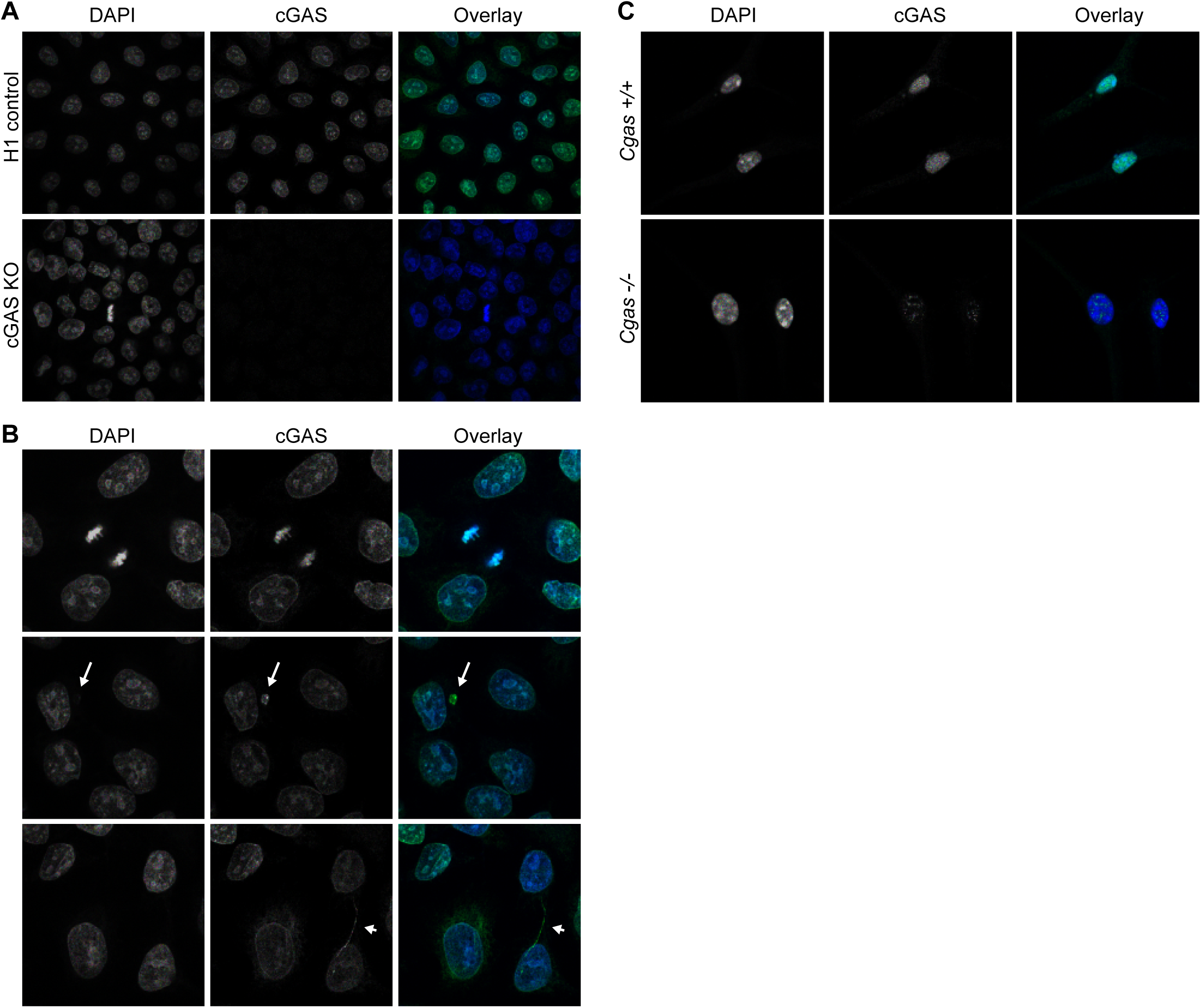
Endogenous cGAS is predominantly a nuclear protein. (**A**) Clonal lines of control HeLa cells (top row) and cGAS KO HeLa cells were fixed with methanol, stained with an antibody to human cGAS, counter-stained with DAPI, and visualized by confocal microscopy. (**B**) We noted three reproducible patterns of cGAS localization in addition to the nucleus: condensed mitotic chromatin (top row), structures resembling micronuclei (middle row), and tendril-like bridges between cells. (**C**) Mouse *Cgas*+/+ and *Cgas*−/− primary bone marrow-derived macro-phages were stained using a mouse antibody to cGAS and processed as in (A).

To extend our findings to primary cells of another species, we searched for antibodies that could identify endogenous mouse cGAS by microscopy. Using primary bone marrow-derived macrophages from wild-type and cGAS-deficient mice and a mouse-specific cGAS antibody, we found nearly exclusive localization of mouse cGAS to the nucleus (Figure 1C). However, even with optimization of blocking conditions and antibody dilutions, we noted that cGAS-deficient mouse macrophages displayed a pattern of nuclear staining that was distinct in its distribution and less abundant than the cGAS staining of wild-type cells (Figure 1C). Despite the imperfect background fluorescence, this was the most sensitive and specific cGAS staining we could identify among the antibodies that we tested. These data reveal that, contrary to expectation, cGAS is primarily a nuclear protein in both human and mouse cells.

### cGAS is tethered tightly in the nucleus

We sought to reconcile the nuclear localization of endogenous cGAS in Figure 1 with the widely accepted notion that cGAS is primarily a cytosolic protein. To track endogenous cGAS localization thoroughly, we modified a protocol for salt-based elution of histones from purified nuclei (Shechter et al., 2007). We prepared extracts separating cytosol from nuclei using a solution containing 0.2% NP-40 detergent followed by low speed centrifugation. After washing the pellets with detergent-free lysis buffer, we lysed the nuclei in a solution of 3mM EDTA and 0.3mM EGTA in water. Following this zero salt nuclear lysis, the pellets remaining after centrifugation were treated with stepwise increases of NaCl in a buffer containing 50 mM Tris-HCl pH 8.0 and 0.05% NP-40. We tracked endogenous cGAS throughout this sequential extraction and elution protocol using eight different cell lines from humans and mice, sampling primary cells, immortalized cells, and tumor cells, as well as rapidly dividing cells and non-dividing cells (differentiated mouse macrophages). We monitored the specificity of the extractions using the cytosolic protein tubulin, the nuclear zero salt lysis-enriched protein LSD1, and nuclear pellet-localized histones H2B and H3. In all of these cells, we found that the vast majority of cGAS was not only in the nuclear fractions, but it was remarkably resistant to salt-based elution (Figure 2A). In most of these cells, a NaCl concentration of 0.75M or higher was required to solubilize the majority of cGAS, similar to the amount of salt required to initiate the liberation of histones (Figure 2A). Importantly, the salt elutions reflect sequential treatments of the same nuclear fractions such that the sum of all the cGAS signals in these fractions can be compared to the cytosolic extracts in order to determine the relative amounts of cGAS in the cytosol and nucleus. These comparisons, calculated by densitometry analysis (Figure 2B), corroborate the microscopy studies in Figure 1 and reveal that the great majority of endogenous cGAS is in the nucleus, not in the cytosol. Moreover, our findings demonstrate that the conventional nuclear washes of ~420mM NaCl that are typically used to isolate cGAS are insufficient to liberate cGAS from the nucleus (Sun et al., 2013). Such tight tethering of cGAS in the nucleus cannot be explained by its low intrinsic affinity for DNA, the dissociation constant of which has been estimated at 1-2 μM (Civril et al., 2013; Li et al., 2013).

**Figure 2:**
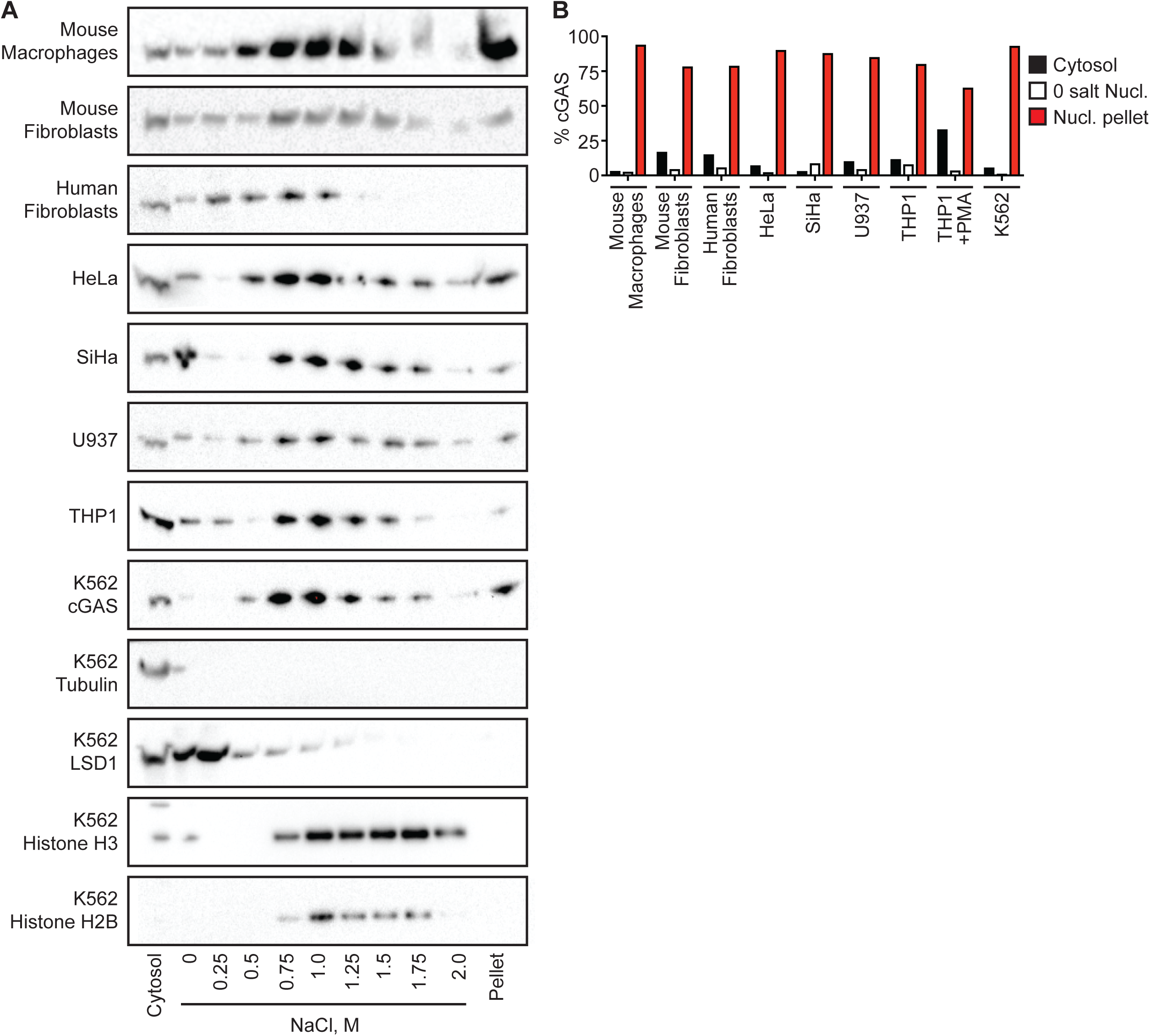
cGAS is tightly tethered in the nucleus. (**A**) Eight mouse and human cell lines were separated into cytosol and nuclear fractions, followed by sequential stepwise elutions of nuclear pellets with the indicated concentrations of NaCl. cGAS and the indicated control proteins were monitored throughout the elution by western blot. (**B**) Densitometry measurements quantitating the relative amounts of endogenous cGAS protein in the cytosol, the 0 salt nuclear lysis, and the cumulative nuclear pellet.

### cGAS is nuclear regardless of cell cycle phase or activation state

One potential explanation for the nuclear localization of cGAS, particularly in dividing cells like HeLa cells, is that this reflects the observed association of cGAS with condensed mitotic chromatin. Thus, recently divided cells might still retain cGAS in the nucleus before its redistribution to the cytosol via mechanisms that remain unexplained. Importantly, the fact that almost 95% of cGAS is nuclear in largely post-mitotic, differentiated primary mouse macrophages argues against this possibility (Figures 1C and 2). We further tested this by tracking the localization of cGAS throughout controlled cell cycles in K562 cells and HeLa cells. We first treated K562 cells with the well-characterized cyclin-dependent kinase 4/6 (CDK4/6) inhibitor PD 0332991, which arrests cells early in the G1 phase of the cell cycle (Toogood et al., 2005). Propidium iodide (PI) staining of DNA in these cells revealed that the great majority of these cells remained arrested in G1 for 72 hours after treatment, with a small residual population arrested in G2 and a depletion of cells in S-phase (Figure 3A). We sampled the localization of endogenous cGAS each day for three days after treatment. Based on our observation that nuclear cGAS is resistant to salt elution up to 0.75M (Figure 2), we used a widely used commercial extraction kit to separate cytosol from the nuclear proteins that elute in ~420 mM NaCl (nuclear supernatant, NS), and we additionally examined the residual pellets to visualize the entire pool of tightly tethered cGAS (nuclear pellet, NP). Despite the arrest of these cells for three days, we observed no change in the distribution of endogenous cGAS, which remained tightly tethered in the nuclear pellets (Figure 3B). We then used double-thymidine block to synchronize HeLa cells at the G1/S boundary (Bootsma et al., 1964), followed by release that resulted in a uniform progression through a single cell cycle. At 4 and 8 hours post release, PI staining confirmed uniform populations of cells in S and G2/M phases, respectively (Figure 3C). By 24 hours, the cells had become asynchronous again (Figure 3C). At all of these time points, we found that the localization of the majority of cGAS to the nuclear pellet did not change (Figure 3D). Thus, endogenous cGAS is a tightly tethered nuclear protein, regardless of cell cycle phase.

**Figure 3:**
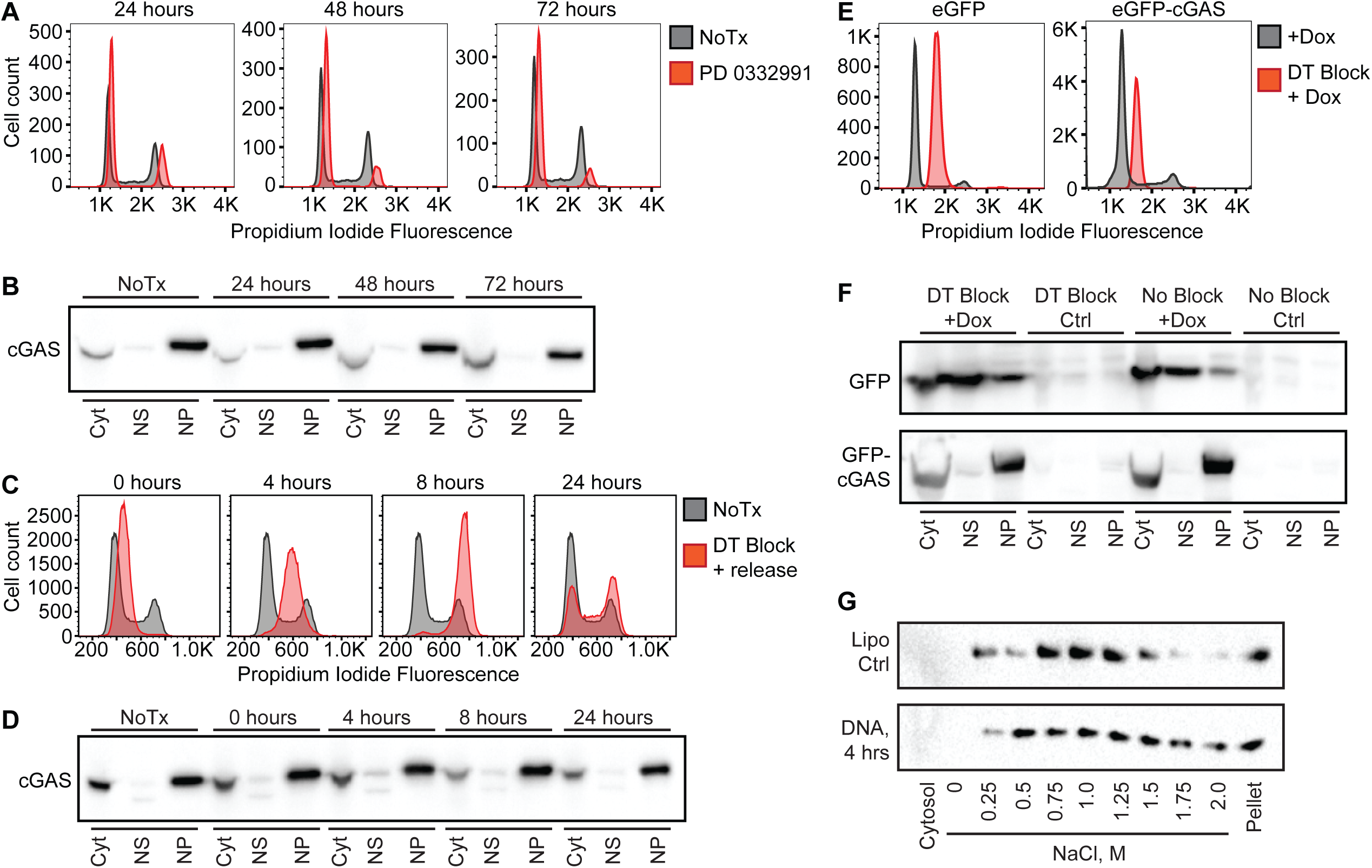
cGAS nuclear localization is independent of cell cycle phase or activation status. (**A**) K562 cells were treated with the CDK4/6 inhibitor PD 0332991 and harvested at the indicated time points. Propidium iodide fluorescence was used to measure DNA content by flow cytometry. (**B**) K562 cells treated as described above were separated into cytosol (Cyt), nuclear supernatant (NS), and nuclear pellet (NP), followed by western blotting for endogenous cGAS. (**C**) HeLa cells were arrested at the G1/S border using double thymidine (DT) block, followed by release and harvest at the indicated time points for measurement of DNA content. (**D**) Cells from (C) were fractionated and cGAS localization was determined by western blot. (**E**) cGAS KO HeLa cells were reconstituted with dox-inducible lentivirus encoding GFP alone or GFP-cGAS, induced with 0.1 μg/ml Dox alone, or subjected to DT block and treated with 0.1 μg/ml Dox for the last 16 hours before harvest, fractionation, flow cytometry analysis for DNA content. (**F**) Cells treated as in (E) were analyzed by western blot using anti-GFP antibody. (**G**) HeLa cells were transfected with Lipofectamine alone (Lipo) or with CT-DNA for 4 hours, followed by extraction, salt elution, and western blot for endogenous cGAS.

Next, we tested whether nuclear localization and tethering of cGAS required cell division, given that current models envision cGAS associating with mitotic chromatin during mitotic prometaphase, after dissolution of the nuclear envelope. To do this, we cloned a GFP-cGAS expression construct into a doxycycline-regulated lentivirus vector that enabled transduction of cGAS-deficient HeLa cells followed by selection for these transduced cells in the absence of cGAS expression. Induction of GFP-cGAS expression with doxycycline allowed us to examine its localization in the absence of exogenous DNA stimulation, unlike standard transient transfections in which the plasmid DNA encoding cGAS also serves as a potent activating ligand. We arrested these cells at the G1/S border using a double-thymidine block, and we induced GFP-cGAS expression with doxycycline for the final 16 hours of the treatment, confirming the arrest of the cells at the G1/S border (Figure 3E). We found that the distribution of the induced GFP-cGAS was identical in cycling and arrested cells, with the majority tightly tethered in the nucleus (Figure 3F). These data demonstrate that cGAS is actively imported into the intact nucleus, rather than passively associated with chromatin during mitosis.

Lastly, we asked whether cGAS localization is dependent on its activation state. We transfected HeLa cells with calf thymus DNA and harvested them four hours later, at a time when they were making large amounts of cGAMP (Figure 1 – Figure Supplement 1A). We performed sequential extractions and salt elutions, comparing cGAS distribution in control and stimulated cells. We did not observe any concerted relocalization of cGAS into the cytosol, despite its robust activation at this time point (Figure 3G). These data demonstrate that activation of cGAS does not result in a dramatic redistribution to the cytosol.

### The N-terminus of cGAS is dispensable for nuclear tethering

We next determined the domains of cGAS that contribute to its tight tethering in the nucleus. The core of human cGAS is comprised of a bilobed structure bridged by an alpha-helical spine (Figure 4A; (Civril et al., 2013; Li et al., 2013). In addition, the N-terminal ~150 residues of cGAS form an unstructured domain that is positively charged and refractory to crystallization. Interestingly, this N-terminus of cGAS was recently demonstrated to be essential for its activation by DNA through a process of phase condensation that assembles cGAS on long double-stranded DNA (Du and Chen, 2018). We reconstituted cGAS-deficient HeLa cells with GFP-cGAS fusions of full-length human cGAS and several truncation mutants corresponding to the structural domains of cGAS. Our panel of truncation mutants revealed a number of important features of its nuclear tethering. First, we found that the GFP-cGAS(161-522) truncation mutant lacking the N-terminus remained tethered in the nuclear pellet, and that the isolated N-terminus of cGAS (1-161) primarily localized to the cytosol and nuclear supernatant (Figure 4B). Second, we found that removal of either the alpha-helical spine (161-213) or the C-terminal lobe of cGAS (383-522) resulted in a protein that was mislocalized and, in the case of the 213-522 mutant, also unstable (Figure 4B). Thus, the intact core of cGAS is required for its nuclear tethering.

**Figure 4:**
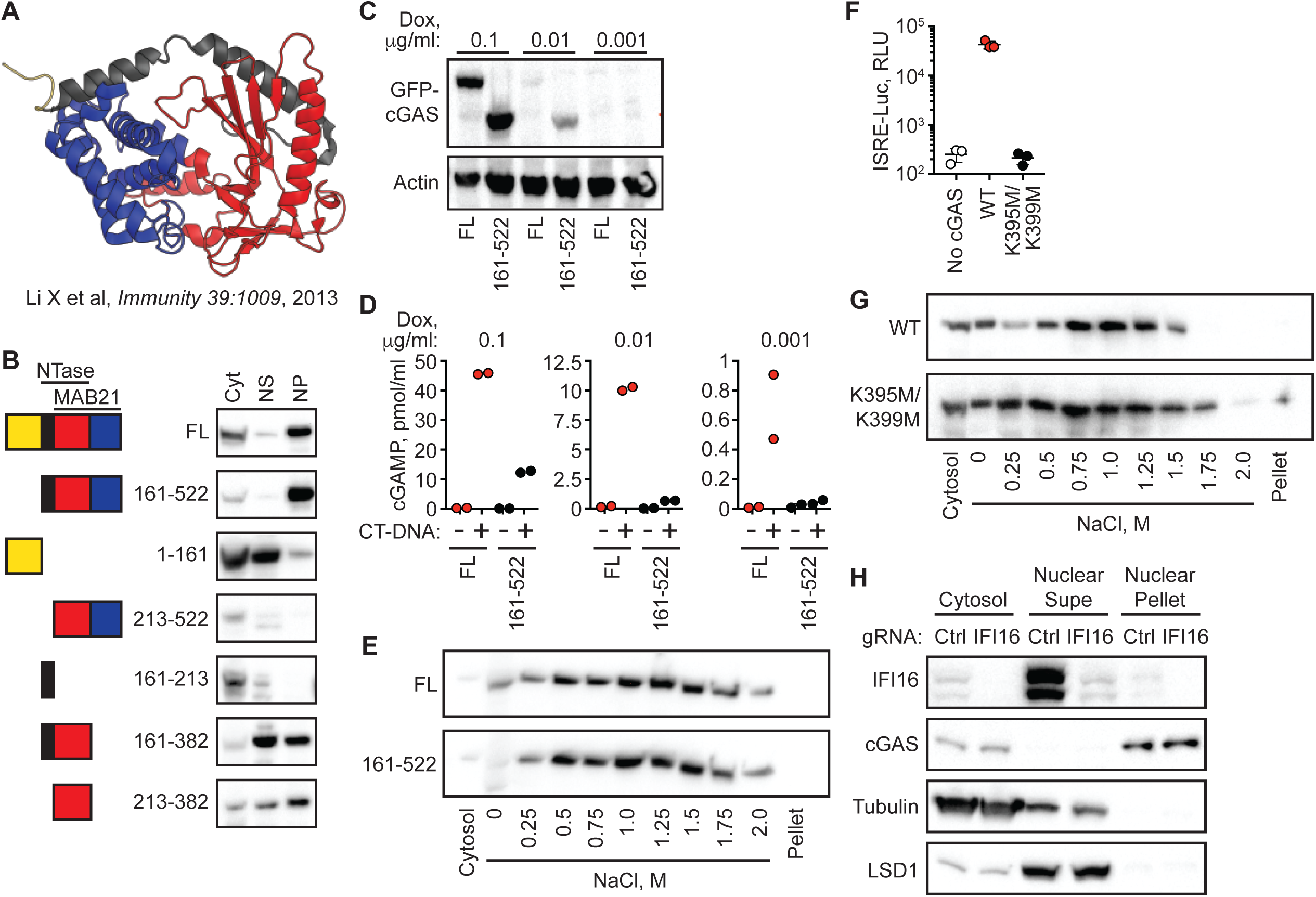
The cGAS N-terminus is dispensable for nuclear tethering. **(A)** Structure of human cGAS with domains colorized. **(B)** TERT-immortalized human foreskin fibroblasts were reconstituted with the indicated Dox-inducible GFP-cGAS lentivirus constructs, treated with 0.1 μg/ml Dox for 24 hours, and then separated into cytosolic (Cyt), nuclear supernatant (NS), and nuclear pellet (NP) fractions. **(C)** cGAS-deficient HeLa cells reconstituted with the indicated GFP-cGAS constructs were induced for 24 hours with three doses of Dox. Whole cell lysates that recover all cGAS were prepared and blotted with anti-GFP antibody. **(D)** Cells from (C) were transfected with CT-DNA for four hours, followed by cGAMP measurement in lysates by modified ELISA. **(E)** Cells described in A-C were treated with 0.1 μg/ml Dox for 24 hours to induce GFP-cGAS expression, then harvested and used for sequential fractionation and salt elution as in Figure 2. **(F)** The indicated mouse cGAS constructs were transiently transfected into HEK293T cells together with STING expression vector and ISRE-luciferase reporter. Luciferase activity in extracts was measured 24 hours later. **(G)** Dox-inducible mouse cGAS constructs were introduced into hTERT-immortalized human fibroblasts. Cells were treated with 0.1 μg/ml Dox for 24 hours followed by fractionation. **(H)** TERT-immortalized human fibroblasts were transduced with lentiCRISPR constructs encoding either control gRNA or gRNA targeting *IFI16,* selected with puromycin to enrich for transduced cells, and then fractionated and blotted with the indicated antibodies.

We compared the full-length and N-terminal deletion mutant of cGAS in more detail. We tested a 100-fold range of dox concentrations that induced varying levels of the GFP-cGAS fusion constructs, from robust to nearly undetectable (Figure 4C). Consistent with the recent definition of the requirement of the N-terminus for cGAS condensation onto DNA (Du and Chen, 2018), we found that the mutant lacking the N-terminus of cGAS was severely compromised for DNA-activated cGAMP production at all levels of expression when compared to full-length cGAS (Figure 4D). However, sequential salt elution of the two forms of cGAS revealed nearly identical distribution and similarly tight tethering in the nucleus (Figure 4E). These data reveal two important points about the requirements for cGAS localization and its activation. First, the domains that tether cGAS in the nucleus are distinct from the domain required for efficient DNA-induced activation, revealing separate mechanistic processes that govern the resting and activated states of cGAS. Second, it could be argued that our conditions of cGAS extraction and salt elution might result in the unnatural condensation of cytosolic cGAS onto DNA that becomes liberated during the extraction, which could then cause it to co-sediment with nuclei during the low speed centrifugation step. Our findings argue against this possibility because the N-terminal domain that is essential for cGAS condensation is dispensable for cGAS nuclear tethering.

We tested two additional potential cGAS tethering mechanisms. First, we found that a K395M/399M mutant of mouse cGAS that is unable to be activated by DNA was still strongly localized in the salt-resistant fractions of the nucleus (Figure 4F, 4G; (Civril et al., 2013). Second, we used a validated lentiCRISPR approach to disrupt the gene encoding IFI16 (Gray et al., 2016), which has been proposed to interact with cGAS and contribute to its activation (Orzalli et al., 2015). We found that IFI16 was extracted by ~420 mM salt into the nuclear supernatant and that IFI16 disruption resulted in no change in cGAS protein levels or its tight tethering in the nuclear pellet (Figure 4H).

### Intact chromatin is required for cGAS tethering

cGAS binds to DNA in a sequence-independent fashion, and its association with chromatin is thought to be generic, mediated by its intrinsic affinity for DNA. To broadly assess the requirement for chromatin in the tethering of nuclear cGAS, we treated nuclei with two different broad-spectrum nucleases that digest both DNA and RNA. First, we digested the nuclear extracts after the zero salt lysis step for 30 minutes at 37 degrees C with micrococcal nuclease, which was sufficient to convert the vast majority of chromatin into nucleosome-protected DNA fragments under 200 bp in length (Figure 5A). Second, we added salt-active nuclease (SAN) to both the 250 mM and 500 mM salt elution steps, which eliminated all detectable DNA from the samples (Figure 5A). In both cases, the nuclease digestions resulted in a collapsed salt elution profile for histones and cGAS, with the majority of cGAS released from the pellets at 250 mM salt (Figure 5B). These data demonstrate that intact chromatin is required for the organization of cGAS nuclear tethering.

**Figure 5:**
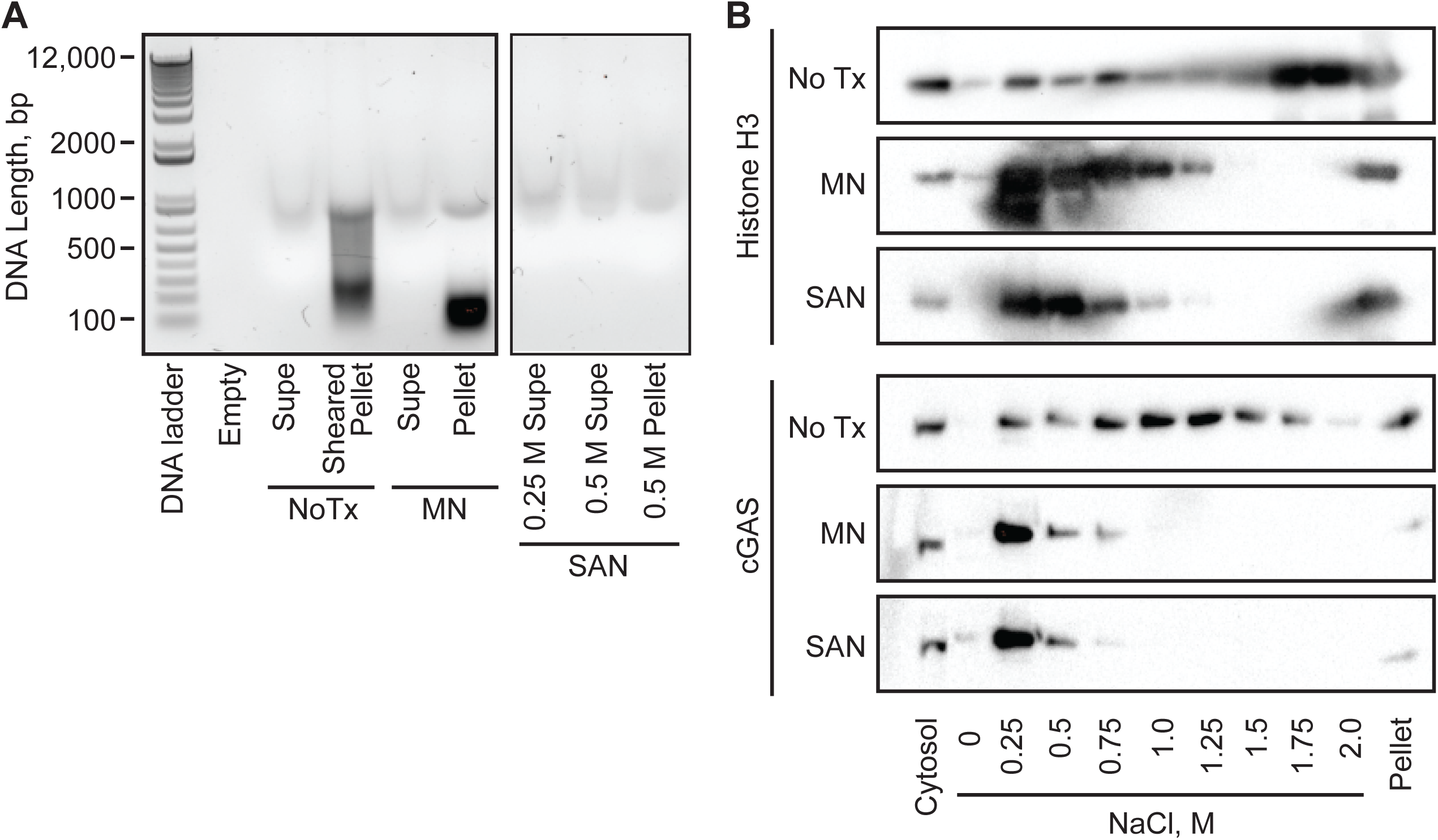
Intact chromatin is required for cGAS tethering. **(A)** K562 cell nuclear extracts were left untreated (NoTx), treated after the 0 salt wash step with micrococcal nuclease (MN), or treated at both the 0.25M and 0.5M NaCl elution steps with Salt Active Nuclease (SAN). Supernatants and pellets were collected, and the untreated pellet was sonicated to shear large genomic DNA. DNA was extracted, run on an agarose gel, and visualized with SYBR Safe. (**B**) Extracts treated as described above were used for sequential salt elution, followed by blotting for Histone H3 and cGAS.

## Discussion

Put simply, the cytosolic DNA sensing model holds that cGAS is a cytosolic protein that is activated by binding to double-stranded DNA that appears in the cytosol (Stetson and Medzhitov, 2006). Our findings suggest that DNA detection by cGAS is more complex than this simple model, and they warrant further study of the relationship between resting and activated cGAS, together with the spatial and biochemical transitions that accompany its activation.

We show, using microscopy and biochemical fractionation, that the great majority of endogenous cGAS is nuclear prior to its activation, in all cells tested, in all phases of the cell cycle, and independent of mitosis. Many recent studies, most using overexpressed, tagged cGAS, have shown images of cGAS localized to the cytosol and absent from the nucleus, although nuclear cGAS has been observed in mouse fibroblasts expressing GFP-cGAS (Yang et al., 2017) and in mouse hematopoietic stem cells (Xia et al., 2018). Others have recently reported regulated translocation of a fraction of overexpressed cGAS from the cytosol to the nucleus in response to DNA damage (Liu et al., 2018). Finally, studies of cGAS localization to micronuclei envision the recruitment of cytosolic cGAS to cytosolic micronuclei upon rupture of the membrane surrounding these structures (Bartsch et al., 2017; Dou et al., 2017; Gluck et al., 2017; Harding et al., 2017; Mackenzie et al., 2017; Yang et al., 2017). We cannot explain these disparate findings, but our data demonstrate that endogenous cGAS is in the nucleus prior to its activation, in the absence of exogenous DNA damaging agents, and prior to the appearance of micronuclei that require mitosis for their formation.

We found that endogenous cGAS is tethered tightly in the nucleus by a force that is remarkably resistant to salt extraction, which explains why this pool of cGAS has been missed when conventional cytosolic and nuclear extracts have been reported in prior studies. We demonstrate that the N-terminal 160 amino acids of cGAS, which are essential for its potent activation by DNA, are dispensable for this nuclear tethering. Finally, we show that cGAS nuclear tethering requires intact chromatin. Our findings must be reconciled with two important characteristics of cGAS. First, the known intrinsic affinity of cGAS for DNA is weak (Kd ~1-2 μM) and cannot alone account for the tight tethering that we observe (Civril et al., 2013; Li et al., 2013). Second, cGAS is not constitutively active, even though we show that the great majority of cGAS is nuclear and that its tethering requires chromatin. We therefore speculate that DNA is not the primary tether of cGAS. Instead, we propose that some protein component(s) of higher order chromatin organize cGAS tethering and regulate its access to DNA, thus preventing its activation by self DNA. If DNA itself were the cGAS tether, it would be difficult to rationalize how cGAS that is in constant contact with billions of base pairs of genomic DNA would be competent to respond to foreign DNA that would be of much lower abundance in a DNA virus-infected cell.

What is the nature of the compartment occupied by cGAS? Our future studies are aimed at defining its constituents and their role in regulation of cGAS access to DNA. A recent study identified the nuclear protein NONO as an important factor for localization of a pool of cGAS to the nuclear compartment that is extractable by ~400 mM salt (Lahaye et al., 2018). It will be interesting to explore the role of NONO and other candidate tethering factors in the regulation of the highly salt-resistant pool that constitutes the majority of nuclear cGAS. More broadly, we propose that this compartment might be a platform to regulate transactions between other DNA-binding proteins and DNA in the nucleus, serving as a mechanism to prevent inappropriate activation of enzymes that require and/or act on DNA. The term “nuclear matrix” was coined decades ago to account for the organization of proteins and macromolecules that are not in direct contact with DNA (Berezney and Coffey, 1974; Nickerson, 2001). While this term remains controversial because of lack of agreement on what precisely constitutes the nuclear matrix, it was proposed to be largely sequestered from DNA, and that specific DNA sequences could be enriched in this compartment after viral infection or plasmid transfection (Jankelevich et al., 1992; Jones and Su, 1987). We wonder if nuclear cGAS might occupy a compartment in the nucleus analogous to the matrix, thus regulating its access to DNA and preventing its constitutive activation by the genome.

Which pool of cGAS – cytosolic or nuclear – is competent to respond to DNA? It could be that the tightly tethered pool of nuclear cGAS is permanently inactivated, and that the small fraction of cytosolic cGAS is primarily responsible for cGAMP production upon activation by DNA. However, the fact that the great majority of resting cGAS is in the nucleus must be considered in future studies on its activation. We note that the production of the diffusible second messenger cGAMP by cGAS provides a simple explanation for how activated nuclear cGAS could trigger the cytosolic signaling complex of STING, TBK1, and IRF3.

We propose a model of “regulated desequestration” of nuclear cGAS and DNA that is required for its full activation, which is then followed by the phase separation and condensation of cGAS onto DNA that was recently and compellingly demonstrated (Du and Chen, 2018). Although this phase condensation model was proposed to explain activation of cGAS in the cytosol, it could also explain how nuclear cGAS becomes activated, especially since we show that the N-terminus that is essential for DNA-induced condensation is dispensable for nuclear localization and tethering. Thus, two separate processes govern the resting and activated states of cGAS, and we propose that a deeper understanding of these will illuminate new aspects of cGAS biology, with implications for mechanisms of self/non-self discrimination by the innate immune system.

## Materials and Methods

### Immunofluorescence Microscopy

HeLa cells or primary mouse bone marrow-derived macrophages (BMMs) were seeded onto glass coverslips overnight, then fixed and permeabilized with ice cold methanol at -20 degrees C for 10 minutes. Cells were then washed in PBS and blocked at room temperature for 2 hours (HeLa cells Roche block in PBS; BMM in 5% normal goat serum in PBS). Cells were then incubated with primary antibody in block over night at 4 degrees C (HeLa cell CST D1D3G Ab at 1:50; BMM CST D3080 Ab at 1:250). Cells were washed in PBS and incubated with secondary Ab (goat anti-rabbit Alexafluor 488, Invitrogen) at 1/500 for 1 hour at room temperature. Cells were then washed with PBS, stained with DAPI and mounted on glass slides with ProLong Gold Antifade Mountant (Thermo Fisher). Images were captured with a Nikon C2RSi Scanning Laser Microscope, using a Plan Apo VC 60× Oil DIC N2 objective in the 405 and 488 dichroic channels with NISElements Software, and then pseudocolored using Fiji open source software.

### Generation of cGAS knockout HeLa cells

LentiCRISPR vector generation and lentiviral transductions were done as described previously (Gray et al., 2016). Clonal lines of HeLa cells were generated by limiting dilution and then assessed for targeting by Sanger sequencing, western blot analysis, and functional assays for cGAMP production. The guide RNA H1 off-target control is 5’-(G)ACGGAGGCTAAGCGTCGCAA (Sanjana et al., 2014), where the (G) denotes a nucleotide added to enable robust transcription from the U6 promoter; and cGAS 5’ - GGCGCCCCTGGCATTCCGTG<u>CGG</u>, where the underlined sequence denotes the Protospacer Adjacent Motif (PAM).

### Salt Extractions

We modified a published protocol for histone extraction (Shechter et al., 2007). Cells were pelleted, washed in PBS, resuspended in 1ml extraction buffer (10mM Hepes pH 7.9, 10mM KCl, 1.5mM MgCl_2_, 0.34M sucrose, 10% glycerol, 0.2% NP-40, and Pierce protease inhibitors), and incubated on ice for 10 minutes with occasional vortexing. Nuclei were spun at 6,500 × g for 5 minutes at 4 degrees C. The cytosolic fraction (supernatant) was collected for further analysis. Nuclei were then washed for 1 minute on ice in extraction buffer without NP-40 and spun at 6,500 × g for 5 minutes at 4 degrees C. Pelleted nuclei were then resuspended in 1 ml zero salt buffer (3mM EDTA, 0.2 mM EGTA, and protease inhbitors), and vortexed intermittently for 1 minute (10 seconds on, 10 seconds off). Nuclei were then incubated on ice for 30 minutes, vortexing for 15 seconds every 10 minutes. Lysates were then spun at 6,500 × g for 5 minutes at 4 degrees C. The zero salt supernatant was collected for further analysis. The remaining pellets were then resuspended in first salt buffer (50 mM Tris-HCl, pH 8.0 0.05% NP-40, 250mM NaCl), incubated on ice for 15 minutes with vortexing for 15 seconds every 5 minutes. Lysates were spun at full speed (15,000 rpm) at 4 degree C for 5 minutes. Supernatants were collected for further analysis. Subsequent salt extractions were performed on the pellet with sequential increase in NaCl concentration (500mM, 750mM, 1M, 1.25M, 1.5M, 1.75M, and 2M). Samples in each salt wash were incubated on ice for 15 minutes with vortexing for 15 seconds every 5 minutes. Supernatants following each salt condition were collected for further analysis. The final pellet was then resuspended in salt buffer with 2M NaCl and sonicated with a Covaris M220 focused ultrasonicator at 5% ChIP (factory setting), or digested with Salt Active Nuclease (SAN) where the buffer was supplemented with 20mM MgCl_2_. All samples were supplemented with denaturing SDS-PAGE sample buffer, separated on acrylamide gels, transferred to membranes for western blot (0.2 μM pore size for histone blots, 0.45 μM pore size for all other blots), and blotted with the indicated primary and secondary antibodies using standard approaches. Western blot images were acquired and densitometry analysis was performed using a BioRad Chemidoc and associated software.

### NE-PERS kit modification

The NE-PERS kit instructions (Thermo Fisher) were followed completely, with the following modification: after spinning the pellet out of the NER buffer, the supernatant was removed saved as “nuclear supernatant (NS)”. The remaining pellet was resuspended in a volume of NER buffer equal to the first, and either sonicated (using Covaris M220 5% ChIP factory setting), or digested with SAN in NER buffer supplemented with 20mM MgCl_2_. This was then saved as “nuclear pellet (NP)”.

### PD 0332991 treatment

K562 cells were treated with 8 μM of PD-0332991 (Selleckchem) in complete media. At the indicated time points cells were harvested for analysis by western blot or for flow cytometry. For flow cytometry analysis, cells were washed in PBS, resuspended in 70% ethanol, and then placed at -20 degrees C. Cells were then washed in PBS, resuspended in PI staining buffer (40μg/ml Propridium iodide; 20μg/ml RNaseA in PBS) and then incubated at 37 deg C for 30 minutes before analysis on a Becton Dickinson LSRII flow cytometer.

### Double Thymidine block

Cells were seeded onto plates to achieve 40% confluency. The next day cells were treated with 2mM thymidine in complete media for 19 hours. Cells were then washed in warm PBS and rested in complete media for 9 hours. Cells were then treated again with 2mM thymidine in complete media for 16 hours. Cells in which GFP-cGAS expression was induced were treated with 1 μg/mL doxycycline for the duration of the second thymidine treatment. Cells were then either harvested for analysis by western blot or flow cytometry; or washed and returned to complete media and harvested at post-release time points. Flow cytometry analysis was performed as described above.

### cGAMP quantitation assay

Cells were plated at 100,000 cells/well in a 24 well tissue culture dish. 24 hrs later, cells were transfected with either 10 μg/ml CT-DNA in lipofectamine 2000 (Invitrogen; ratio of 1 μl lipofectamine per 1 μg CT-DNA; (Stetson and Medzhitov, 2006), or with an identical volume of lipofectamine 2000 alone. 4 hours later, cells were harvested and lysates were prepared using cGAMP EIA assay protocol provided by manufacturer (Arbor Assays), in a volume of 200 μl sample suspension buffer.

### Constucts

The pSLIK-Neo doxycycline-inducible lentiviral vector was obtained from Addgene and modified to replace the Neo cassette with a blasticidin resistance cassette. GFP fusions to the murine cGAS open reading frame were generated by PCR mutagenesis and designed to incorporate a four-glycine flexible linker between the last amino acid of GFP and the first amino acid of cGAS. Lentivirus production and blasticidin selection were done using standard techniques.

### Doxycycline-inducible cGAS constructs

cGAS-deficient HeLa cells were reconstituted with lentiviruses encoding GFP or the indicated GFP-murine cGAS fusions. Cells were plated at 50,000 cells per well in a 24 well plate for quantitation of cGAMP, or 250,000 cells per well in a 6-well plate for salt extractions. 24 hours later, cells were treated with doxycycline for 24 hours. Then, cells were harvested directly for anti-GFP WB from the 6-well dishes, and the 24-well dishes were transfected with either 10 μg/ml CT-DNA in lipofectamine 2000, or with lipofectamine 2000 alone. 4 hours later, lysates were prepared and analyzed for cGAMP content as described above (whole cells lysed in 200 μl sample suspension buffer).

### Nuclease digestions and salt elutions

Salt extractions were performed as described above with the following modifications. 1×10^6^ cells were used for each condition. Following the zero salt wash, all samples were resuspended in digestion buffer, (50 mM Tris pH 8.0, .05% NP-40, 1 mM MgCl2, 5mM CaCl_2_). For the MNase digestion, MNase was added at 20,000 gel units per sample. For the SAN digestion, 50 units SAN nuclease plus 20 mM MgCl_2_ were added at both the 250 and 500 mM NaCl steps All samples were incubated at 37 degrees C for 10 minutes. Samples were then spun down and supernatants and pellets were separated and then processed for western blots using the salt elution protocol. For analysis of DNA content, the MN-digested supernatants and pellets were analyzed after digestion and before commencement of salt elution. For the SAN digestion, the pellet was collected after the 500mM NaCl elution for assessment of DNA content. DNA was run on an agarose gel, stained using SYBR-Safe reagent (Apex Bio), and visualized using a BioRad Chemidoc.

### ISRE luciferase assay

HEK 293T cells were seeded at 25,000 cells per well in a 96 well plate. They were then transfected with the following plasmids: 20ng of hSTING-pCDNA3 (Lau et al., 2015), 5ng of pISRE-luciferase (Clontech), 60ng of empty pCDNA3, and 5ng of cGAS-pCDNA3 (Lau et al., 2015). Eighteen hours later, cells were lysed in passive lysis buffer (Promega). Lysates were assayed on a Centro LB 960 luminometer (Berthold) using luciferase reagent (Promega E4550) according to the manufacturers protocol.

### Experimental replicates and reproducibility

All data presented in this paper are representative of 2-4 independent experiments with comparable results.

## Acknowledgements

We are grateful to Quinton Dowling and Neil King for their help with cGAS structural analysis, to Andrew Oberst and Naeha Subramanian for the pSLIK-Neo vector, to Michael Gale, Jr for use of the confocal microscope, and to the entire Stetson lab for helpful discussions.

---

**Figure 1, Figure Supplement 1.**
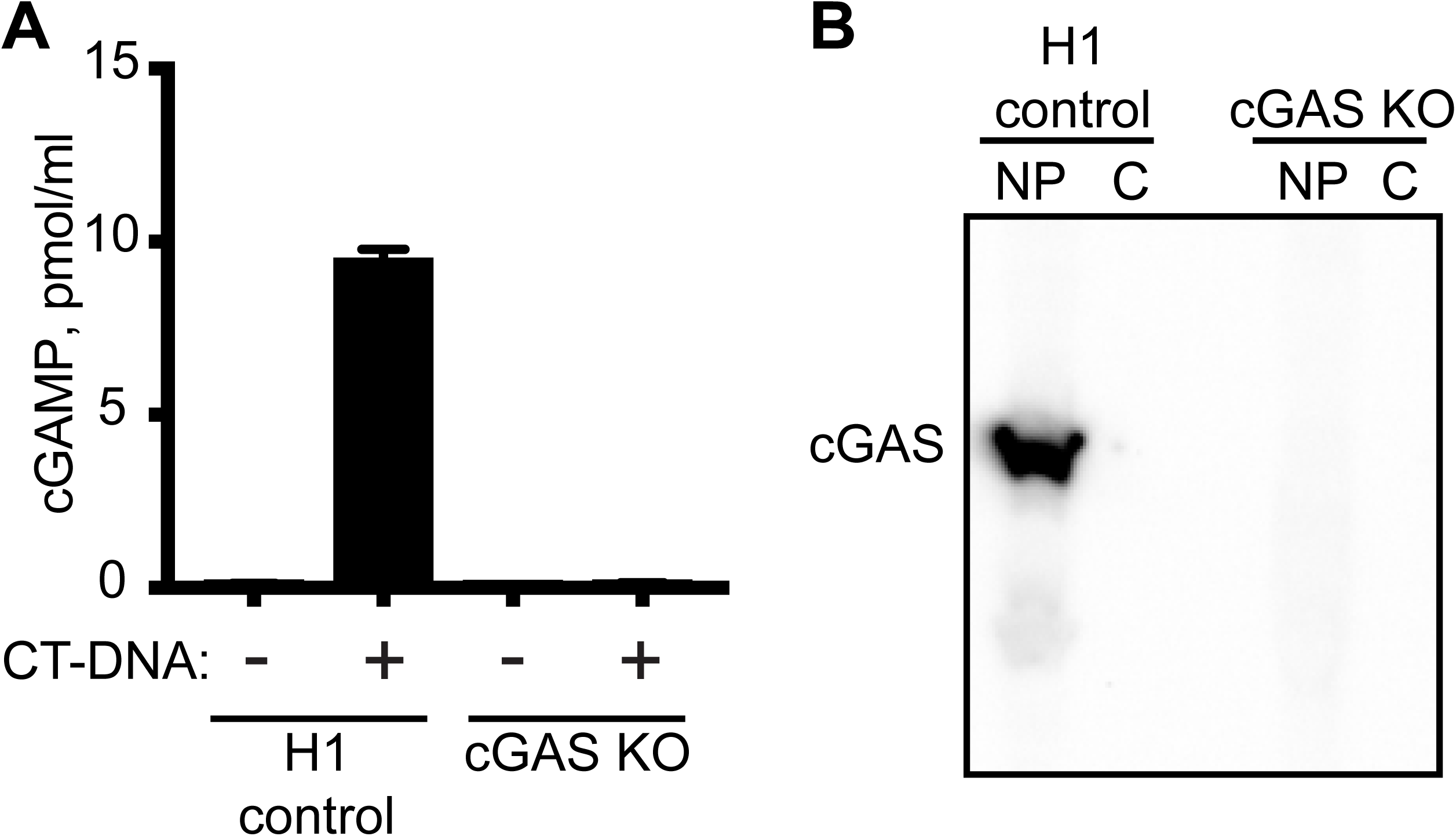
Characterization of clonal cGAS KO HeLa cells. (**A**) HeLa cells were transduced with LentiCRISPR encoding control or cGAS-targeted gRNAs, cloned by limiting dilution, and tested for production of cGAMP in cell lysates four hours after transfection of calf thymus DNA (CT-DNA). (**B**) Lysates from control and cGAS KO clonal HeLa cells were separated into cytosol (C) and nuclear pellet (NP), and then blotted for endogenous cGAS.

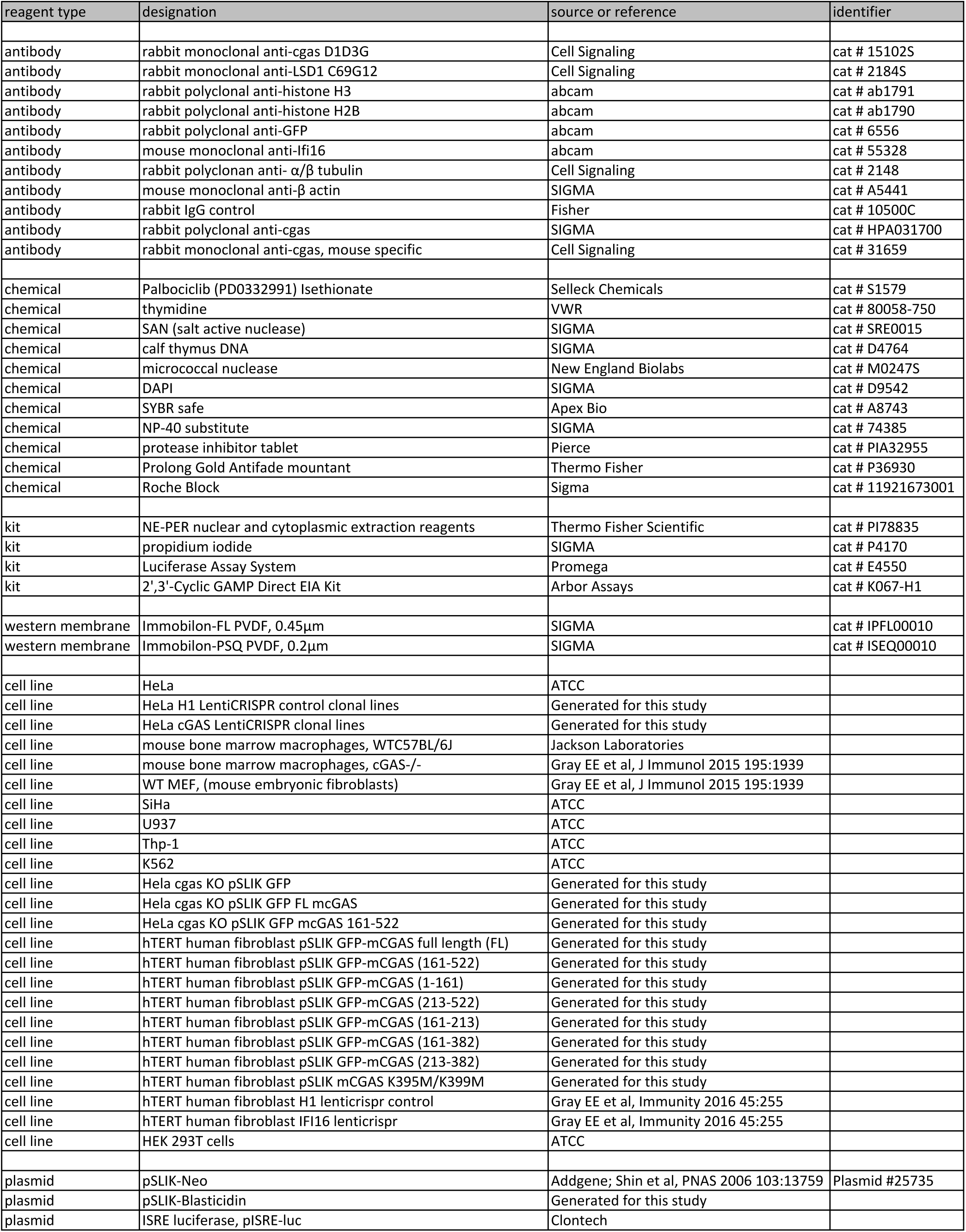

